# Adaptive Integration of Heterogeneous Foundation Models to Find Histologically Predictable Genes in Breast Cancer

**DOI:** 10.64898/2026.04.05.716435

**Authors:** Hao Nguyen, Chaoyi Li, Can Peng, Peter Simpson, Nan Ye, Quan Nguyen

## Abstract

Foundation models for computational pathology have rapidly emerged as powerful tools for extracting rich biological and morphological representations from histopathology images. However, variations in model architecture, pre-training data, and optimization objectives often lead to task-dependent performance, rather than universal generalization. As a result, effective strategies for integrating their complementary strengths are essential to fully realize the potential of foundation models for robust histopathology analysis. Meanwhile, recent breakthroughs such as spatial transcriptomics provide an unprecedented opportunity to integrate genetic and histopathology information from the same patient sample, thereby maximizing both molecular and anatomical pathology insights. Specifically, each model’s embedding is first mapped to gene-level predictions via a dedicated prediction head, enabling model-specific feature utilization. A lightweight weighting network then adaptively aggregates these predictions to produce a unified and robust output at gene and spatial location levels. Across multiple spatial transcriptomics datasets, our approach consistently outperforms both individual foundation models and classical ensembling methods. Focusing on breast cancer, we observe substantial gains in prediction accuracy for clinically relevant PAM50 subtype markers and drug-target genes. Moreover, the proposed framework improves interpretability by revealing model-specific contributions and specialization at the gene level. Overall, our work presents an effective solution to integrating multiple foundation models for enhancing the genetic analyses of histopathology images.

## 1 Introduction

In recent years, self-supervised learning based pathology foundation models (FMs) have substantially advanced computational pathology (CPath), accelerating the translation from research to clinical applications [2]. These models generate rich visual embeddings that capture tissue morphology, thereby improving performance across a wide range of downstream tasks [9, 17]. One important application is spatial gene expression prediction, which infers spatial transcriptomics (ST) measurements from histopathology images. Although an increasing number of publicly available FMs have been developed at different scales using diverse architectures, learning objectives, and datasets, most existing approaches for gene expression prediction still rely on a single FM. In contrast, biological modalities often encode complementary information, suggesting that a single-model approach may not fully exploit the diverse representations learned under different training paradigms [2, 1]. Recent benchmarking studies [3, 5, 15, 13] further demonstrate that no single FM consistently outperforms others across all genes or tissue types. This observation indicates that different FMs capture distinct and partially non-overlapping biological signals. Therefore, effectively integrating multiple FMs to leverage their complementary strengths is a promising direction for improving gene expression prediction.

While several ensemble methods have been explored, existing strategies typically produce a single globally fused representation, implicitly assuming that one shared representation is optimal for all tasks [15]. However, in ST, different genes may depend on distinct morphological cues and contextual patterns. By relying on a global representation, current ensemble methods fail to account for such gene-specific heterogeneity. To address this limitation, we propose a gene- and spot-specific adaptive ensemble framework that dynamically weights model contributions at both the spatial and gene level. Rather than enforcing a single fused embedding, our method allows model selection to vary across genes and tissue locations, enabling fine-grained specialization for spatial gene expression prediction.

## 2 Related Work

### Foundation Models in Computational Pathology

Pathology FMs have rapidly evolved in scale, data diversity, and clinical applicability. Recent models, including CONCH [10], CHIEF [21], and Virchow2 [24], leverage large-scale histopathology pretraining and incorporate weak supervision, multimodal alignment, and stain robustness across large-scale of whole slide images (WSIs) datasets, thereby improving anatomical coverage and clinical relevance. Distillation-based approaches, such as GPFM [12] and H0-mini [6], further compress knowledge from one or multiple expert models to enhance efficiency and generalization. Table 1 summarizes the details of these FMs. Despite these advances in scale and design, current pathology FMs are not explicitly optimized for fine-grained biological inference. Their embeddings and pretraining objectives may not fully capture the combinatorial and gene-specific morphological patterns required for spatial gene expression prediction [19]. This limitation motivates strategies that can better exploit complementary representations across models.

**Table 1:** FMs evaluated in this study. All models are frozen encoders pre-trained on histopathology images using self-supervised learning.

| Model | Architecture | SSL Method | Training Data (WSI/Tiles) | Params | Embed Dim |
| --- | --- | --- | --- | --- | --- |
| ResNet50 [7] | ResNet-50 | Supervised (ImageNet) | NA/1.2M | 23M | 2048 |
| UNI [4] | ViT-L/16 | DINOv2 | 100K/100M | 307M | 1024 |
| CONCH [10] | ViT-B/16 | iBOT | NA/16M | 86M | 768 |
| GigaPath [22] | ViT-g/14 | DINOv2 | 171K/1.3B | 1.13B | 1536 |
| Virchow2 [24] | ViT-H/14 | DINOv2+ECT+KDE | 3.1M/1.9B | 632M | 1280 |
| CHIEF [21] | CTransPath | Self-distillation | 60K/15M | 304M | 768 |
| Hibou-L [14] | ViT-L/16 | DINOv2 | 1B/1.2B | 307M | 1024 |
| GPFM [12] | ViT-L/16 | MIM+Self-distillation+Expert KD | 72K/190M | 307M | 1024 |
| H0_mini [6] | ViT-B/16 | Distillation | 400K/NA | 86M | 1024 |

### Ensemble Learning

Existing ensemble strategies in computational pathology primarily focus on slide-level classification or global representation learning. Methods such as score averaging, feature concatenation [15], attention-based fusion [11], and uncertainty-aware ensemble [23] have shown improved robustness and performance in classification settings. However, their applicability to **regression tasks for spatial transcriptomics prediction** remains largely unexplored. In particular, existing methods do not explicitly account for gene-specific variability and spatial heterogeneity inherent in this problem. In this work, we extend ensemble learning to this task and introduce an adaptive framework tailored for gene-specific and spatially resolved regression. Our approach enables dynamic model weighting across both genes and spot locations, allowing the ensemble to specialize according to target-specific morphological signals.

## 3 Method

### 3.1 Task Definition

We aimed to predict spatial gene expression from histopathology images, an approach to combine genetics and traditional histology for potential clinical diagnosis applications. Given a histopathology image ℋ paired with *N* spatial locations (spots), and a set of *M* pre-trained pathology FMs 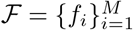, each spot is represented by *M* model-specific embeddings. Our goal is to predict the expression of *G* genes at all *N* locations, producing a matrix **Ŷ** ∈ ℝ^*N*×*G*^ that matches the ground-truth expression matrix **Y** ∈ ℝ^*N*×*G*^.

### 3.2 Feature Extraction

#### Patch Extraction

For each spatial location **s**_*n*_ = (*x*_*n*_, *y*_*n*_), a square image patch is extracted 𝒫_*n*_ from the corresponding H&E image, centered at (*x*_*n*_, *y*_*n*_) with size 224 × 224 pixels at 20× magnification (approximately 112 *µ*m × 112 *µ*m tissue area). This resolution matches the standard input size of most vision backbones and provides sufficient local morphological context around each spot.

#### Foundation model embeddings

Each image patch 𝒫_*n*_ is encoded by the *M* foundation models (we use *M* =9; Table 1) to obtain model-specific embeddings:

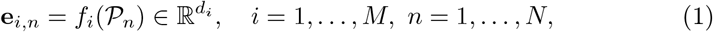

and we stack embeddings across all *N* spots to form:

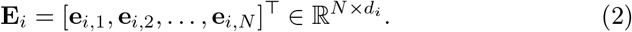

These embeddings serve as inputs to the proposed adaptive ensemble framework.

### 3.3 Adaptive Assemble Approach

Existing ensemble strategies in CPath often produce a single global fused representation. In ST, however, different genes can express differently depend on distinct morphological cues, and the most informative FM may vary across genes and spatial locations. We therefore developed an adaptive ensemble framework that combines (i) model-specific prediction heads and (ii) a learnable weighting module that performs gene- and spot-wise routing, as illustrated in Figure 1.

**Fig. 1:**
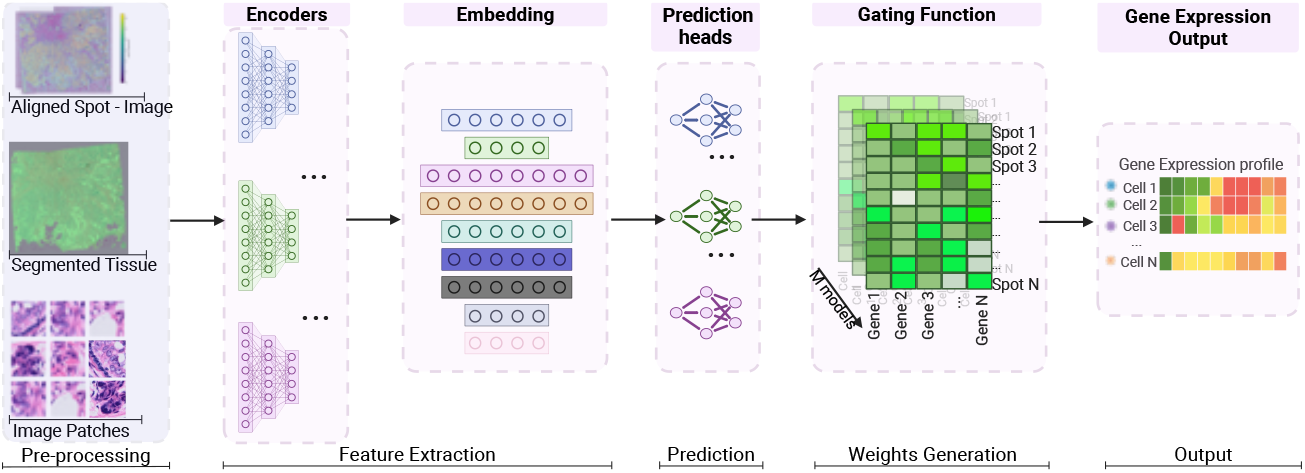
Overview of the proposed framework. ST data on WSI are split into pairs of image tiles and gene expression labels. The model integrates multiple FM embeddings and predicts gene expression at each spatial location by a gating function that computes spot- and gene-aware weighting.

#### Per-model prediction heads

Given heterogeneous spot embeddings 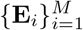 with 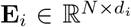 ,we first attach a lightweight, model-specific prediction head to each FM:

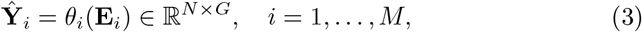

where *θ*_*i*_ is an 3-layer MLP mapping *d*_*i*_ → *h*→ *h/*2 → *G* and produces per-spot predictions for all *G* genes.

#### Adaptive ensembling

The per-model predictions are combined using spot- and gene-aware weighting to predicts weights conditioned on the concatenated embeddings, allowing the selected models to vary by both spot and gene as described below:

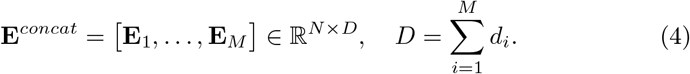

A gating network *ϕ* (2 layer MLP: *D*→*h*^′^ → (*G* × *M*)) outputs spot-dependent weight logits:

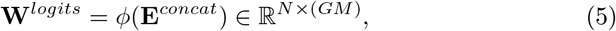

which we reshape into **W**^*logits*^ ∈ ℝ^*N*×*G*×*M*^ and normalize over models:

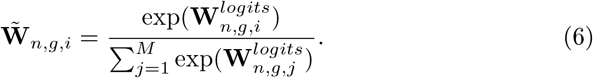

The final prediction is then

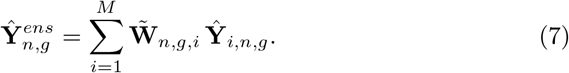

This gating enables spatially adaptive and gene-specific model selection.

#### Training objective

We trained all learnable parameters end-to-end by minimizing the mean squared error between the ensemble prediction and ground-truth expression:

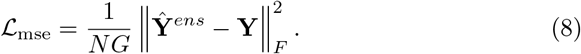

The optimized parameters are 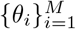 and *ϕ*. We optimized using AdamW with the learning rate of 1 × 10^−4^, a batch size of 512, and trained for 300 epochs.

## 4 Experiments

### 4.1 Datasets

We evaluated 9 pathology FMs (ResNet50, UNI, CONCH, GigaPath, Virchow2, h0mini, CHIEF, GPFM, and Hibou-L) on spatial gene expression prediction using (i) 5 samples of STimage xenium breast cancer dataset (BRCA - Xenium) [18] and (ii) 4 samples of IDC subset of HEST dataset (IDC) [8].

### 4.2 Data Pre-processing

WSIs were acquired at 20× magnification and stored in standard formats (e.g., .tif, .svs). Each WSI is paired with ST data containing gene expression measurements at defined tissue coordinates. ST platforms provide coordinates 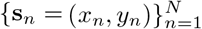 that are registered to the corresponding WSI. Gene expression data **Y**^*raw*^ ∈ ℝ^*N*×*G*^ were processed by log-normalization.

#### Benchmark Pipeline

To ensure a fair comparison with prior work, we adopted the HEST-Bench[8] protocol consisting of four steps: **(1) Feature extraction**: extracting embeddings from 224×224 H&E patches using frozen encoders (batch size 128, 4 workers); **(2) Dimensionality Reduction**: applying PCA to reduce the features to 256 dimensions; **(3) Linear probing**: training a Ridge regression model (*ℓ*_2_ regularization) to predict gene expression levels from the reduced embeddings; **(4)** *k***-fold cross-validation**: aggregating results across predefined splits to compute mean and standard deviation. The primary metric is the Pearson Correlation Coefficient (PCC) between the predicted value and the ground truth, computed per gene by pooling expression values across all spatial locations from all test samples. All experiments were ran on a GPU with a batch size of 512.

### 4.3 Results

#### Overall Performance

Figure 2 shows the per-gene PCC distribution across all methods on both datasets. On the BRCA-Xenium dataset, our ensemble achieves 0.567 ± 0.193, outperforming the strongest single FM, Virchow2 (0.552 ± 0.198), and substantially surpassing the ResNet50 baseline (0.478 ± 0.205). Notably, the ensemble also reduces prediction variability, indicating improved consistency across genes. The gain is more pronounced on IDC, where our method reaches 0.615 ± 0.176, compared to 0.597 ± 0.178 for Virchow2 and 0.503 ± 0.186 for ResNet50, corresponding to a 22.2% relative improvement over the baseline. Across both datasets, the ensemble consistently ranks first, demonstrating robust generalisation and the effectiveness of adaptive model integration.

**Fig. 2:**
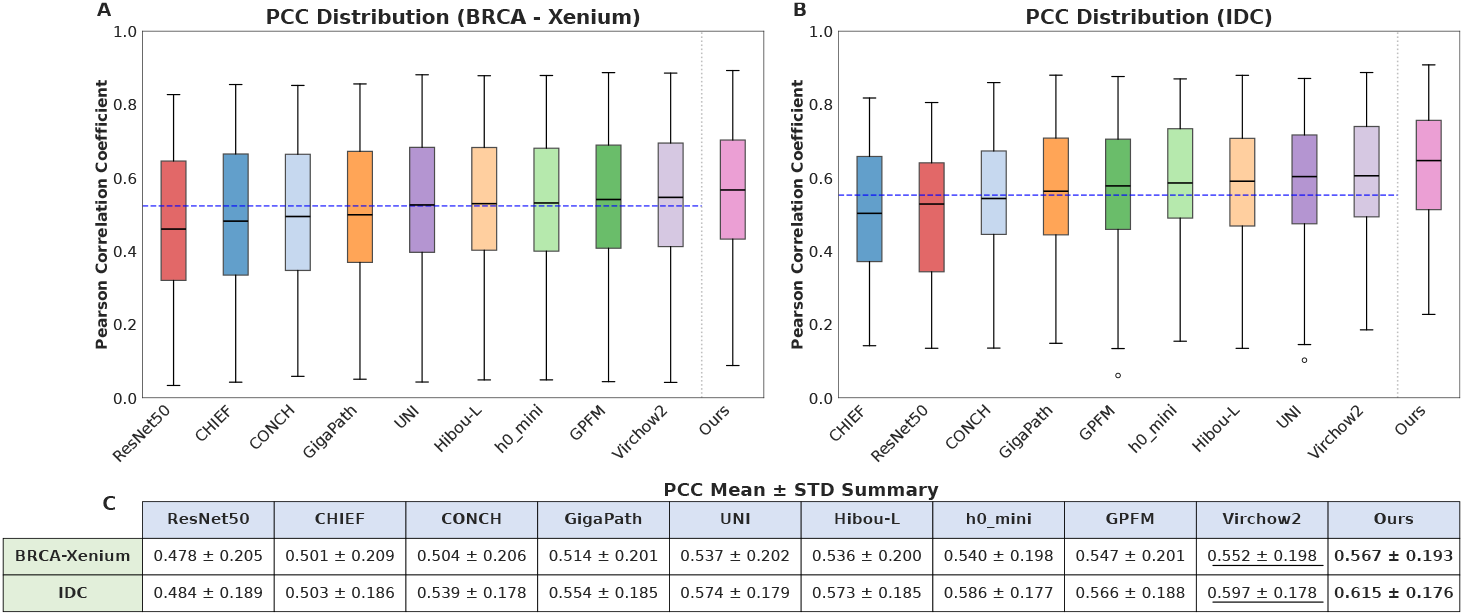
Per-gene PCC distribution across nine FMs and our adaptive ensemble on the BRCA-Xenium dataset. Mean ± standard deviation is reported above each box (for the top 50 highly variable genes of the IDC dataset and all xenium v1 genes panel of the BRCA-Xenium dataset). The dashed blue line marks the average PCC of single encoders. Best is **bold**, second best is underlined.

#### Relevant Gene Markers for Breast Cancer Diagnosis

Figure 3 reports per-gene improvements relative to the ResNet50 baseline for PAM50 subtype markers and additional clinical genes. Our ensemble achieves the highest or joint-highest improvement in nearly every gene category across both datasets. For luminal markers, SFRP1 improves by 70.2% on BRCA-Xenium and 154.9% on IDC; notably, CONCH, h0_mini, Virchow2 underperform ResNet50 on ESR1, underscoring the limit of single-model reliance. Basal markers show the most pronounced gains, EGFR improves by 774.1% on BRCA-Xenium, with KRT5 and KRT14 following similar trends on both datasets, indicating that basal-associated morphology particularly benefits from multi-model integration. For immunotherapy-related genes, our ensemble achieves the largest gains on CD274 (+10.0%) and PDCD1 (+18.7%), whereas several single models underperform the baseline on CD274, suggesting sensitivity to model-specific bias. Adaptive ensemble integration reduces these biases and yields the most robust prediction. Overall, our framework achieves the highest per-gene PCC on 90.7% genes of BRCA-Xenium and 74% genes of IDC, confirming consistent superiority over all individual FM baselines. Collectively, these findings demonstrate that heterogeneous model integration is particularly advantageous for biologically complex and clinically critical gene targets. Consistent with the quantitative gains, Figure 4 further illustrates improved spatial coherence and biological plausibility in the predicted gene expression maps.

**Fig. 3:**
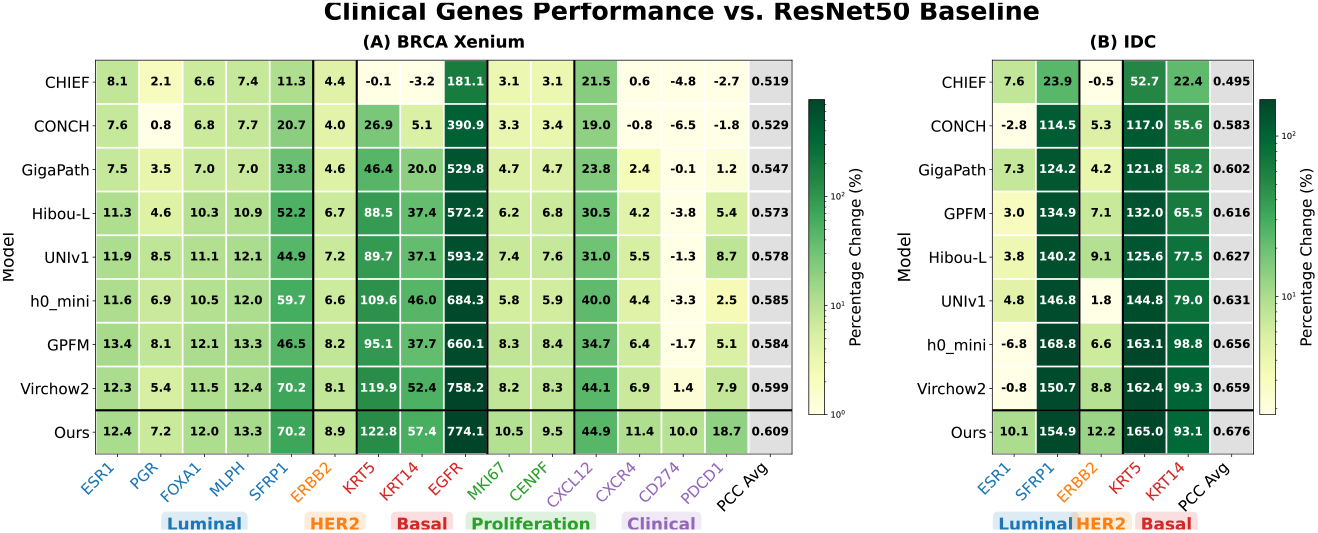
Percentage improvement in PCC over the ResNet50 baseline for clinically relevant gene markers across FMs and our ensemble, on the BRCA-Xenium dataset (A) and IDC dataset (B). Subtypes (Luminal, Basal, Proliferation, HER2 and clinical markers) are indicated by the colored legend beneath the heatmap, and black vertical lines denote boundaries between subtype groups. Colour intensity follows a log scale. Improvements are computed relative to ResNet50: positive values indicate performance gains, negative values indicate regression.

**Fig. 4:**
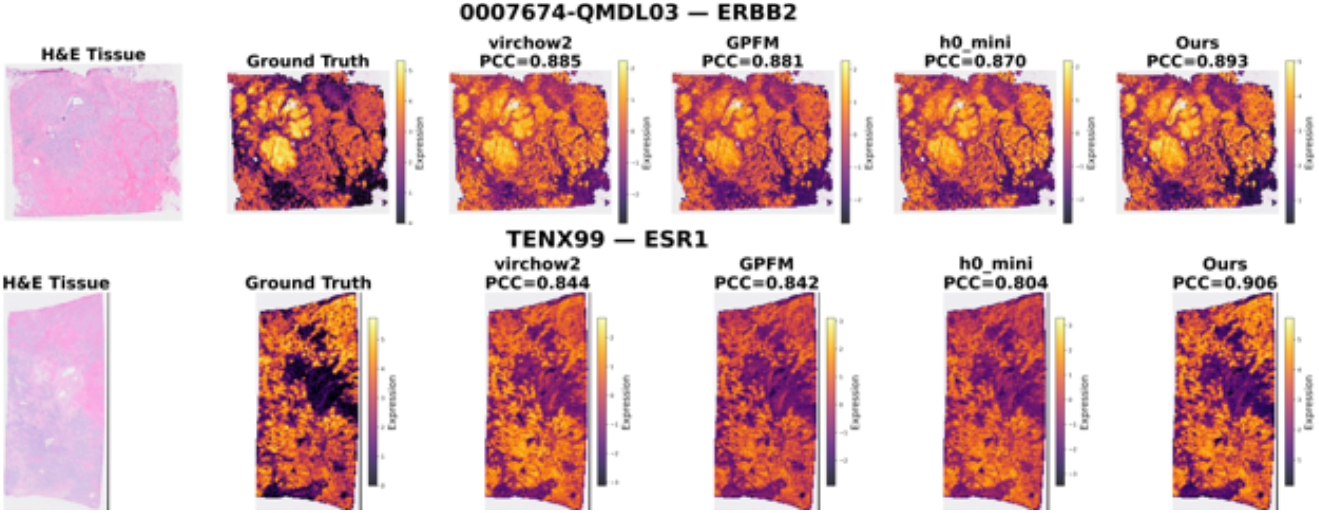
Spatial gene expression prediction maps for ERBB2 (BRCA-Xenium, a single cell ST dataset) and ESR1 (IDC, a lower resolution dataset). From left to right: H&E tissue, ground truth, and predictions from Virchow2, GPFM, h0_mini, and our ensemble.

#### Ablation study

We compared our method against four alternative fusion strategies: (i) *cross-attention* [20], which treats each model as a separate attention head and learns to attend across model outputs; (ii) *gate attention* [16], which employs element-wise gating to modulate model contribution, (iii) *simple concatenation*, which concatenates all embeddings and passes them through a single MLP prediction head, (iv) *unified projection*, which projects all heterogeneous embeddings into a shared dimension before a single prediction head. As shown in Table 2, our adaptive weighting consistently outperforms all alternatives on both datasets, demonstrating that dynamically gating across independently pre-trained FMs is more effective than both global fusion strategies and attention-based aggregation.

**Table 2:** Ablation study comparing fusion strategies across datasets. Mean PCC (↑) ± standard deviation for multiple genes (the performance gain per gene is shown in Fig 3).

|  | Cross-Att | Gate-Att | Concat | Unified-Proj | Ours |
| --- | --- | --- | --- | --- | --- |
| BRCA-Xenium | $0.557 \pm 0.197$ | $0.551 \pm 0.198$ | $0.555 \pm 0.193$ | $0.556 \pm 0.196$ | **$0.567 \pm 0.193$** |
| IDC | $0.587 \pm 0.195$ | $0.595 \pm 0.183$ | $0.610 \pm 0.173$ | $0.612 \pm 0.178$ | **$0.615 \pm 0.176$** |

## 5 Conclusion

In this paper, we present an adaptive ensemble framework for spatial gene expression prediction that integrates heterogeneous pathology foundation models through learnable gene- and spatial-level weighting mechanisms. Across independent breast cancer datasets, our approach consistently outperforms all individual models, with particularly strong gains on clinically relevant PAM50 markers and immunotherapy-related targets. Importantly, the learned weights provide interpretable evidence of gene-specific model specialization, offering possible explanations to link tissue morphology and biological features of breast cancer with performance gain and insights into how different foundation models capture complementary biological signals. Future work will extend this framework to other cancer types and incorporate uncertainty-aware selection strategies to further improve robustness and generalizability.

## References

1. Bahrami, M., Richter, T., Schmacke, N.A., Lavandera, A.E., Theis, F.J.: From modality-specific to compositional foundation models for cell biology. Cell Systems 17(2) (2026)

2. Bilal, M., Raza, M., Altherwy, Y., Alsuhaibani, A., Abduljabbar, A., Almarshad, F., Golding, P., Rajpoot, N.: Foundation models in computational pathology: A review of challenges, opportunities, and impact (2025), 2502.08333

3. Campanella, G., Chen, S., Singh, M., Verma, R., Muehlstedt, S., Zeng, J., Stock,, Croken, M., Veremis, B., Elmas, A., et al.: A clinical benchmark of public self-supervised pathology foundation models. Nature Communications 16(1), 3640 (2025)

4. Chen, R.J., Ding, T., Lu, M.Y., Williamson, D.F.K., Jaume, G., Song, A.H., Chen, B., Zhang, A., Shao, D., Shaban, M., et al.: Towards a general-purpose foundation model for computational pathology. Nature Medicine 30(3), 850–862 (2024)

5. Chen, S., Campanella, G., Elmas, A., Stock, A., Zeng, J., Polydorides, A.D., Schoenfeld, A.J., Huang, K.L., Houldsworth, J., Vanderbilt, C., et al.: Benchmarking embedding aggregation methods in computational pathology: A clinical data perspective (2024), 2407.07841

6. Filiot, A., Dop, N., Tchita, O., Riou, A., Dubois, R., Peeters, T., Valter, D., Scalbert, M., Saillard, C., Robin, G., et al.: Distilling foundation models for robust and efficient models in digital pathology. In: International Conference on Medical Image Computing and Computer-Assisted Intervention (MICCAI). pp. 162–172. Springer (2025)

7. He, K., Zhang, X., Ren, S., Sun, J.: Deep residual learning for image recognition. In: Proceedings of the IEEE Conference on Computer Vision and Pattern Recognition (CVPR). pp. 770–778 (2016)

8. Jaume, G., Doucet, P., Song, A., Lu, M.Y., Almagro Pérez, C., Wagner, S., Vaidya,, Chen, R., Williamson, D., Kim, A., et al.: Hest-1k: A dataset for spatial transcriptomics and histology image analysis. Advances in Neural Information Processing Systems 37, 53798–53833 (2024)

9. Li, D., Wan, G., Wu, X., Wu, X., Nirmal, A.J., Lian, C.G., Sorger, P.K., Semenov, Y.R., Zhao, C.: A survey on computational pathology foundation models: Datasets, adaptation strategies, and evaluation tasks (2025), 2501.15724

10. Lu, M.Y., Chen, B., Williamson, D.F.K., Chen, R.J., Liang, I., Ding, T., Jaume, G., Odintsov, I., Le, L.P., Gerber, G., et al.: A visual-language foundation model for computational pathology. Nature Medicine 30(3), 863–874 (2024)

11. Luo, X., Wang, X., Eweje, F., Zhang, X., Yang, S., Quinton, R., Xiang, J., Li, Y., Ji, Y., Li, Z., et al.: Ensemble learning of foundation models for precision oncology (2025), 2508.16085

12. Ma, J., Guo, Z., Zhou, F., Wang, Y., Xu, Y., Li, J., Yan, F., Cai, Y., Zhu, Z., Jin, C., et al.: A generalizable pathology foundation model using a unified knowledge distillation pretraining framework. Nature Biomedical Engineering pp. 1–20 (2025)

13. Mahmood, F.: A benchmarking crisis in biomedical machine learning. Nature Medicine 31(4), 1060–1060 (2025)

14. Nechaev, D., Pchelnikov, A., Ivanova, E.: Hibou: A family of foundational vision transformers for pathology (2024), 2406.05074

15. Neidlinger, P., El Nahhas, O.S.M., Muti, H.S., Lenz, T., Hoffmeister, M., Brenner, H., van Treeck, M., Langer, R., Dislich, B., Behrens, H.M., et al.: Benchmarking foundation models as feature extractors for weakly supervised computational pathology. Nature Biomedical Engineering pp. 1–11 (2025)

16. Qiu, Z., Wang, Z., Zheng, B., Huang, Z., Wen, K., Yang, S., Men, R., Yu, L., Huang, F., Huang, S., et al.: Gated attention for large language models: Nonlinearity, sparsity, and attention-sink-free. arXiv preprint 2505.06708 (2025)

17. Song, A.H., Jaume, G., Williamson, D.F.K., Lu, M.Y., Vaidya, A., Miller, T.R., Mahmood, F.: Artificial intelligence for digital and computational pathology. Nature Reviews Bioengineering 1(12), 930–949 (2023)

18. Tan, X., Mulay, O., Xie, J., MacDonald, S., Kim, T., Zhou, C., Xiong, Z., Tan, S.X., Ye, N., McCart Reed, A., et al.: Robust and interpretable prediction of gene markers and cell types from spatial transcriptomics data. Nature Communications 17(1), 1781 (2026)

19. Tizhoosh, H.R.: Beyond the failures: Rethinking foundation models in pathology. arXiv preprint 2510.23807 (2025)

20. Vaswani, A., Shazeer, N., Parmar, N., Uszkoreit, J., Jones, L., Gomez, A.N., Kaiser, F., Polosukhin, I.: Attention is all you need. Advances in neural information processing systems 30 (2017)

21. Wang, X., Zhao, J., Marostica, E., Yuan, W., Jin, J., Zhang, J., Li, R., Tang, H., Wang, K., Li, Y., et al.: A pathology foundation model for cancer diagnosis and prognosis prediction. Nature 634(8035), 970–978 (2024)

22. Xu, H., Usuyama, N., Bagga, J., Zhang, S., Rao, R., Naumann, T., Wong, C., Gero, Z., González, J., Gu, Y., et al.: A whole-slide foundation model for digital pathology from real-world data. Nature 630(8015), 181–188 (2024)

23. Zhao, J., Lin, S.Y., Attias, R., Mathews, L., Engel, C., Larghero, G., Vremenko, D., Kao, T.W., Lee, T.H., Wang, Y.H., et al.: Uncertainty-aware ensemble of foundation models differentiates glioblastoma from its mimics. Nature Communications 16(1), 8341 (2025)

24. Zimmermann, E., Vorontsov, E., Viret, J., Casson, A., Zelechowski, M., Shaikovski, G., Tenenholtz, N., Hall, J., Klimstra, D., Yousfi, R., et al.: Virchow2: Scaling selfsupervised mixed magnification models in pathology (2024), 2408.00738

